# Large-scale chemical-genetics yields new *Mycobacterium tuberculosis* inhibitor classes

**DOI:** 10.1101/396440

**Authors:** Eachan O. Johnson, Emily LaVerriere, Mary Stanley, Emma Office, Elisabeth Meyer, Tomohiko Kawate, James Gomez, Rebecca E. Audette, Nirmalya Bandyopadhyay, Natalia Betancourt, Kayla Delano, Israel Da Silva, Joshua Davis, Christina Gallo, Michelle Gardner, Aaron Golas, Kristine M. Guinn, Rebecca Korn, Jennifer A. McConnell, Caitlin E. Moss, Kenan C. Murphy, Ray Nietupski, Kadamba G. Papavinasasundaram, Jessica T. Pinkham, Paula A. Pino, Megan K. Proulx, Nadine Ruecker, Naomi Song, Matthew Thompson, Carolina Trujillo, Shoko Wakabayashi, Joshua B. Wallach, Christopher Watson, Thomas R. Ioerger, Eric S. Lander, Brian K. Hubbard, Michael H. Serrano-Wu, Sabine Ehrt, Michael Fitzgerald, Eric J. Rubin, Christopher M. Sassetti, Dirk Schnappinger, Deborah T. Hung

## Abstract

New antibiotics are needed to combat rising resistance, with new *Mycobacterium tuberculosis* (Mtb) drugs of highest priority. Conventional whole-cell and biochemical antibiotic screens have failed. We developed a novel strategy termed PROSPECT (PRimary screening Of Strains to Prioritize Expanded Chemistry and Targets) in which we screen compounds against pools of strains depleted for essential bacterial targets. We engineered strains targeting 474 Mtb essential genes and screened pools of 100-150 strains against activity-enriched and unbiased compounds libraries, measuring > 8.5-million chemical-genetic interactions. Primary screens identified >10-fold more hits than screening wild-type Mtb alone, with chemical-genetic interactions providing immediate, direct target insight. We identified > 40 novel compounds targeting DNA gyrase, cell wall, tryptophan, folate biosynthesis, and RNA polymerase, as well as inhibitors of a novel target EfpA. Chemical optimization yielded EfpA inhibitors with potent wild-type activity, thus demonstrating PROSPECT’s ability to yield inhibitors against novel targets which would have eluded conventional drug discovery.

---

The World Health Organization (1) has declared that antibiotic resistance is one of the greatest threats to human health with tuberculosis (TB) being the deadliest infectious disease, causing more than 1.6 million deaths annually (2). Despite the recent approval of two new drugs (bedaquiline(3) and delamanid (4)),TB drug discovery and development has failed to keep pace with increasing prevalence of multi-, extensively and totally drug resistant TB (5). A fundamental challenge in antibiotic discovery is finding new classes of compounds that kill the causative pathogen, especially by inhibiting novel essential targets. Primary chemical screening using biochemical, target-based assays have yielded compounds lacking whole-cell activity, while conventional whole-cell assays using wild-type bacteria have yielded compounds generally refractory to mechanism of action (MOA) elucidation to enable compound prioritization and progression. The all-too-few successful cases for *Mycobacterium tuberculosis* (Mtb)), the causative agent of TB, illustrate this challenge, with hit compounds found to repeatedly target the same two proteins (MmpL3 and DprE1), leaving the vast majority of Mtb’s∼625 essential proteins unexploited (6-8).

Our goal was to develop a new paradigm of antimicrobial discovery that simultaneously identifies whole cell active compounds and predicts their MOA from primary screening data, thereby incorporating putative target information, instead of simply potency, into hit prioritization. This strategy enables both the exploration of broader target space and discovery of new chemical scaffolds that could not be identified by conventional screening against wild-type bacteria. To enact this strategy, we performed primary chemical screening of hundreds of mutant strains depleted in essential targets (hypomorphs) to generate large-scale chemical-genetic interaction profiles as the output from the primary screen. Although drug hypersensitivity conferred by target depletion is well-established in yeast (9, 10), depleting essential targets in haploid bacteria is challenging. Consequently, chemical screening of hypomorphs in bacteria have been limited either to a single hypomorph of *Staphylococcus aureus* (11) and Mtb (12, 13), or to MOA elucidation of only a single compound against small collections of siRNA-mediated *S*. *aureus* hypomorphs (14, 15); a genome-wide nonessential gene deletion library in *Escherichia coli* has also be used to study single compounds (16). In contrast, our strategy of generating chemical-genetic interaction profiles by screening large hypomorph pools (100-150 hypomorphs) against large chemical libraries (50,000 compounds) – an approach we term PROSPECT (PRimary screening Of Strains to Prioritize Expanded Chemistry and Targets) – dramatically increases detection of active compounds and allows immediate MOA prediction to inform hit prioritization.

PROSPECT yielded ten-fold more hit compounds (> 4000, each associated with chemical-genetic interaction profiles) than conventional whole-cell screening. Using the primary chemical-genetic interaction profiles, we rapidly identified and validated >40 new scaffolds against established targets including DNA gyrase, cell wall biosynthesis, several steps of the folate biosynthesis pathway, tryptophan biosynthesis, and RNA polymerase (RNAP). We identified compounds against novel targets with the discovery of an inhibitor for the uncharacterized, essential efflux pump EfpA. Because hit compounds may have limited activity against wild-type bacteria, we demonstrated the ability to optimize the EfpA inhibitor to druglike potency against wild-type Mtb. PROSPECT can this lead to new molecules against novel targets with potent activity against wild-type bacteria that could not have been found by conventional screening.

## Generating large-scale chemical-genetic interaction profiles from primary screening

We constructed hypomorphs of 474 of the ∼625 essential genes defined in Mtb (17), with each gene target under conditional proteolytic (18, 19) or transcriptional control (Table S1). For proteolytic control, proteins were fused to a carboxterminal DAS-tag, which targets the protein for degradation via SspB-mediated shuttling (Fig. 1a). Because the different essential gene products are naturally expressed at variable levels, each tolerating different degrees of depletion, we constructed up to five hypomorphs, with varying degrees of knockdown, for each gene (Extended Data Fig. 1a-b). Each strain was also engineered to carry a 20-nucleotide genetic barcode flanked by common primer-binding sites for targeted gene identification by multiplexed polymerase chain reaction (PCR) followed by Illumina sequencing. Sequencing read counts of the barcode served as a proxy for individual strain abundance within the pool. In total, we created 2014 strains.

**Figure 1:**
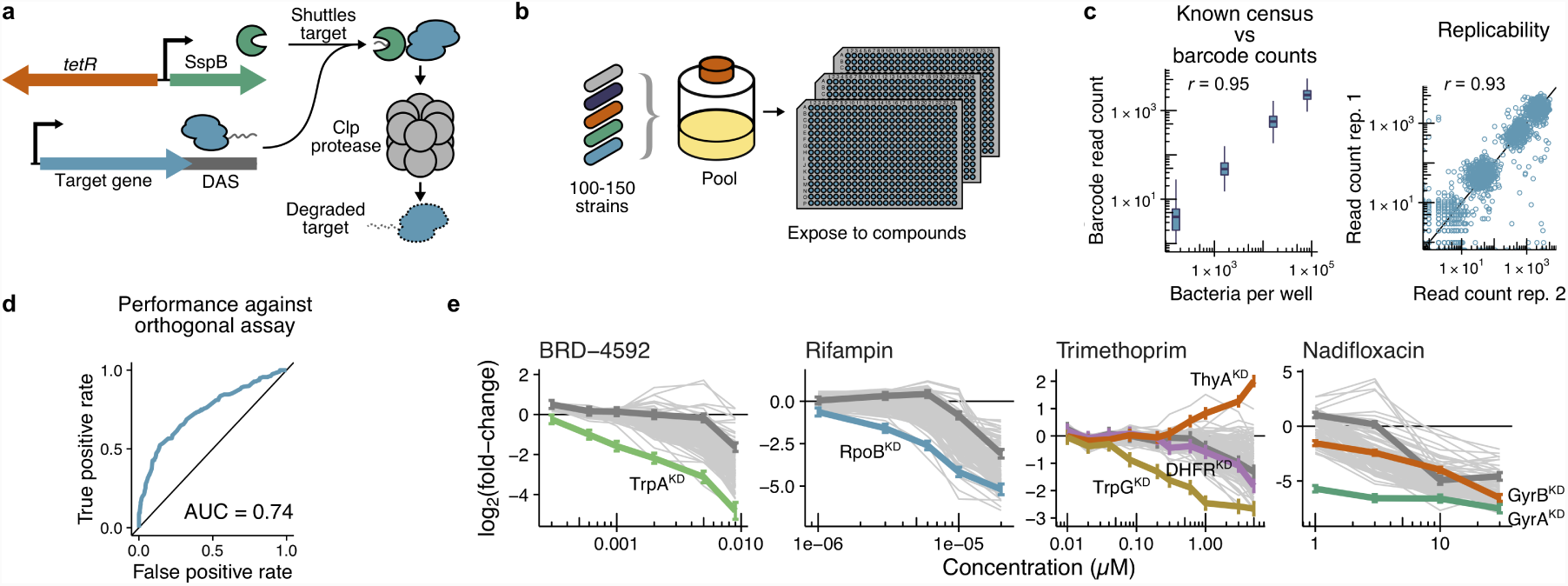
Generating large-scale chemical-genetic interaction profiles from primary screening. **a**, Hypomorph strains were constructed by introducing a DAS-tag at the 3’-end of the gene of interest, with concomitant introduction of a 20-nucleotide barcode and an episomally encoded, regulated SspB gene to control the level of protein depletion. **b**, Barcoded hypomorph strains were pooled and distributed into 384-well plates containing the compound library and incubated for 14 days. **c**, Defined mixtures of barcoded wild type Mtb strains were subjected to census enumeration by sequencing-based barcode counting. The method is accurate across several orders of magnitude with Pearson’s *r* = 0.95 (left panel), and reproducible with Pearson’s *r* = 0.93 between replicates (right panel). **d**, ROC curve showing that primary data was predictive of activity in a confirmatory secondary growth assay. We retested more than 100 compounds predicted to have activity in the primary screen in an orthogonal resazurin-based colorimetric growth assay. Taking 50% inhibition in the secondary assay as ground truth, we demonstrated the primary assay as predictive of real activity that could be detected by more conventional growth methods. The false positive rate is plotted against the true positive rate (blue line); an area under the curve (AUC) more than (black line) indicates a predictor that performs better than chance. **e**, Chemical-genetic interaction profiles showed expected hypersensitivity for compounds of known MOA. Profiles show the LFC (relative to DMSO negative controls) of each strain at each concentration tested (with wild-type Mtb in dark grey and mutants of interest highlighted). Error bars of highlighted strains show 95% confidence interval of the mean. Examples shown are the compound-hypomorph pairs of BRD-4592 with TrpA, rifampin with RpoB, trimethoprim with TrpG, DHFR, and ThyA, and the fluoroquinolone nadifloxacin with GyrA and GyrB.

We established an optimized, multiplexed assay (Fig. 1b) to measure the abundance of each strain in a pool through barcode amplification and sequencing (Extended Data Fig. 1c) (20-23). By mixing strains at known abundances spanning three orders of magnitude, we confirmed that barcode counts were an accurate proxy for strain abundance (*r* = 0.93 for log-transformed barcode count replicates; *r* = 0.95 for known cell abundance and barcode counts) (Fig. 1c). We then used this assay to determine the growth rate of each engineered strain in a pool containing 100-150 strains, including barcoded wild-type in 384-well plates over two weeks. For inclusion in the final screening pool used for all subsequent multiplexed experiments, we selected a single mutant corresponding to each essential gene with the most degradation while maintaining growth similar to the wild-type strain (Extended Data Fig. 2a). Using rifampin (RIF) as a positive control, we found excellent assay performance across the dynamic range for all strains (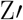 factors > 0.5; Extended Data Fig. 2b).

Finally, we developed a barcode counting (ConCensusMap), and inference (ConCensusGLM) computational pipeline, which provided a log2(fold-change) (LFC) of strain abundance upon compound exposure compared to DMSO control and associated p-value. The LFC vectors across all strains for a compound is termed the chemical-genetic interaction profile.

## Chemical-genetic interactions of Mtb-bioactive compounds

We initially screened a library of 3226 small-molecules enriched for compounds with activity against wild-type Mtb based on literature reports (Extended Data Fig. 2c; see Methods). To confirm the reported Mtb activity, we screened the library against GFP-expressing wild-type Mtb and found that 1312 (45%) indeed had an MIC_90_ < 64 µM (Extended Data Fig. 2d). We then screened this chemical library with a pool of 100 Mtb hypomorph strains in duplicate (log-transformed Pearson’s *r* = 0.93) at compound concentrations of 1.1, 3.3, 10, and 30 µM (chosen based on measured MIC_90_ values for the entire library).

In total we measured 1,290,400 chemical-genetic interactions (3226 compounds × 100 strains × 4 concentrations) with the majority (927,025, 71%) being inhibitory (LFC < 0) (Extended Data Fig. 2e). Of these, 55,508 interactions (6%), representing 940 compounds (29%), were strong (*p* < 10^−10^). In a minority of cases, protein depletion conferred resistance to inhibitors of wild-type Mtb; for example, depletion of the mycothiol biosynthesis pathway enzyme cysteine ligase (MshC) resulted in resistance to TB drugs isoniazid (INH) and ethionamide (ETH), which are known to inhibit the enoyl-[acylcarrier-protein] reductase (InhA) (24).

Using an orthogonal growth assay, we retested 112 of the identified hits with evidence of specificity for a subset of strains (*p* < 10^−10^; specificity defined as activity against < 10 strains) to confirm their activities against their respective, predicted hypomorph interactor, wild-type Mtb, and several other hypomorph strains as negative controls. Because growth rate measurements by these different assays cannot be directly compared, we constructed a receiver operating characteristic (ROC) curve to determine how well inhibitory activity in the multiplexed assay predicted activity in the orthogonal growth assay. The ROC area under the curve (AUC) was 0.73 (Fig. 1d), indicating a high true positive rate in the primary assay with a well-controlled false positive rate. Given the complexity of the primary screen, we were reassured that 1375 (52%) of the 2664 strong interactions were confirmed in the secondary assay.

We readily identified interactions between well-characterized inhibitors and hypomorphs corresponding to known targets (Fig. 1e), including between the fluoroquinolones and the DNA gyrase α-subunit (GyrA), RIF and the RNAP β-subunit of (RpoB), and BRD-4592 (25) and the tryptophan synthase α-subunit (TrpA). Interestingly, trimethoprim (TMP), a folate biosynthesis inhibitor known to target dihydrofolate reductase (DHFR), demonstrated a clear interaction with the folate pathway enzyme glutamine amidotransferase (TrpG) rather than DHFR. The ThyA hypomorph also showed resistance to trimethoprim; in the setting of ThyA loss of function, DHFR is not essential (26). Reducing the 400-dimensional interaction profiles (100 strains × 4 concentrations) for each compound to two dimensions using a *t*-distributed stochastic neighbor embedding (t-SNE) (27), we found that compounds known to have the same MOA clustered together, independent of their chemical structures (Fig. 2).

**Figure 2:**
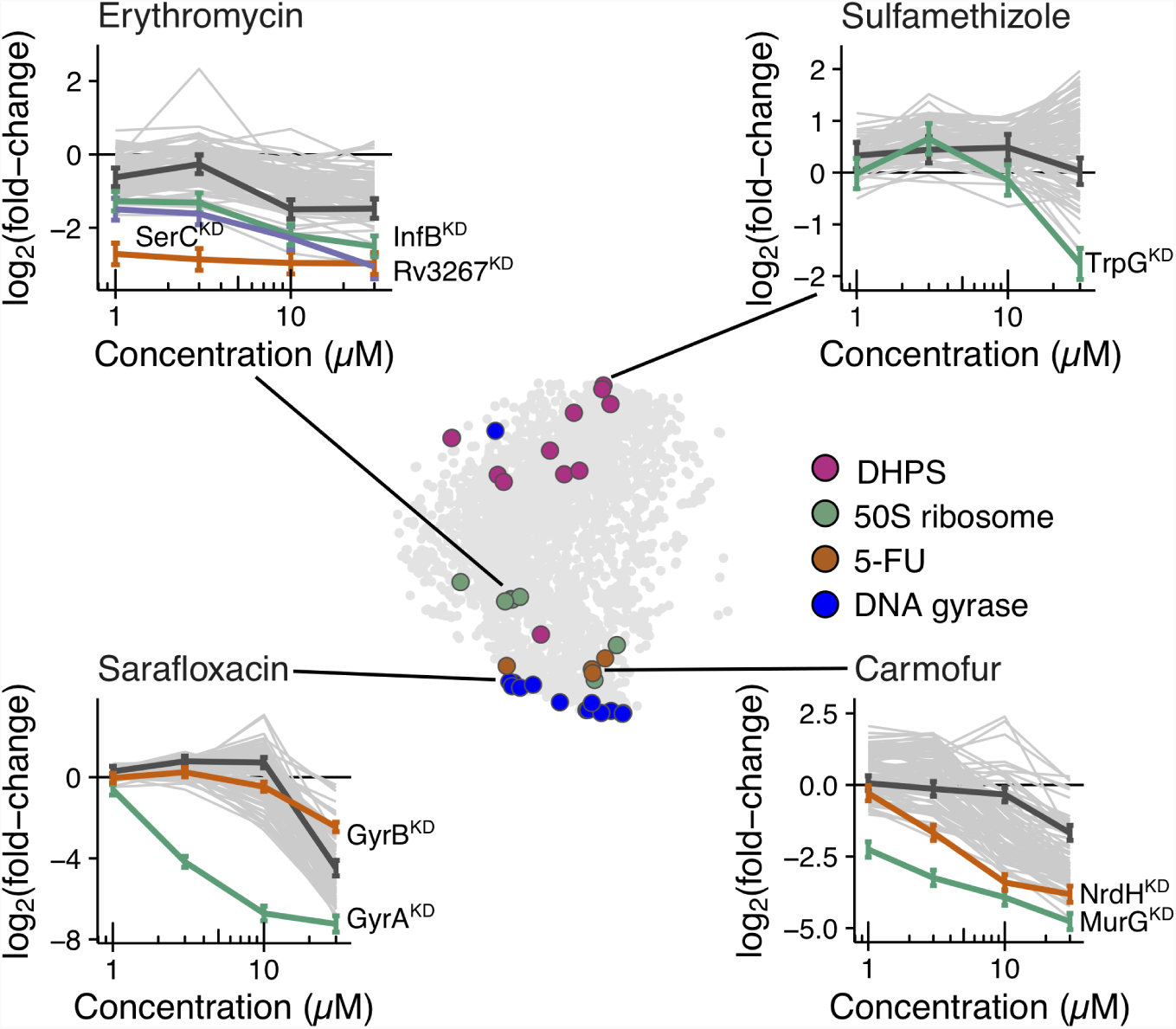
Chemical-genetic interactions of bioactive compounds with known MOA. Using t-SNE to visualize the 400-dimensional dataset reveals the MOA-based clustering of compounds. Four exemplary MOAs are illustrated. Grey circles represent compounds, and colored circles represent a subset of compounds with known MOA. Representative chemical-genetic interaction profiles are shown as in Fig. 1e for compounds representing each of the four exemplary MOAs.

## Discovering new inhibitor classes of well-validated targets using reference data

We identified new compound scaffolds that inhibit well validated, clinical targets based solely on the primary screening data, relative to known reference compounds. Using a core ground truth training set of 107 chemical-genetic interaction profiles for known antimicrobials (Table S4), we trained Lasso classification models (28) – a supervised machine learning method – to identify 39 new inhibitors of DNA gyrase (training set *n* = 14), mycolic acid synthesis (*n* = 6), folate biosynthesis (*n* = 12), and tryptophan biosynthesis.

### DNA gyrase inhibitors

We trained on 14 chemical-genetic interaction profiles of fluoroquinolones, known DNA gyrase inhibitors. Our model’s regression weights suggested that the single most discriminatory feature for gyrase inhibition was strongly decreased fitness of the GyrA hypomorph, which we termed a sentinel strain for this pathway (Extended Data Fig. 3a). The model predicted 55 non-quinolone DNA gyrase inhibitors (Fig. 3a), including novobiocin, a structurally-distinct inhibitor of the DNA gyrase β-subunit (GyrB).

**Figure 3:**
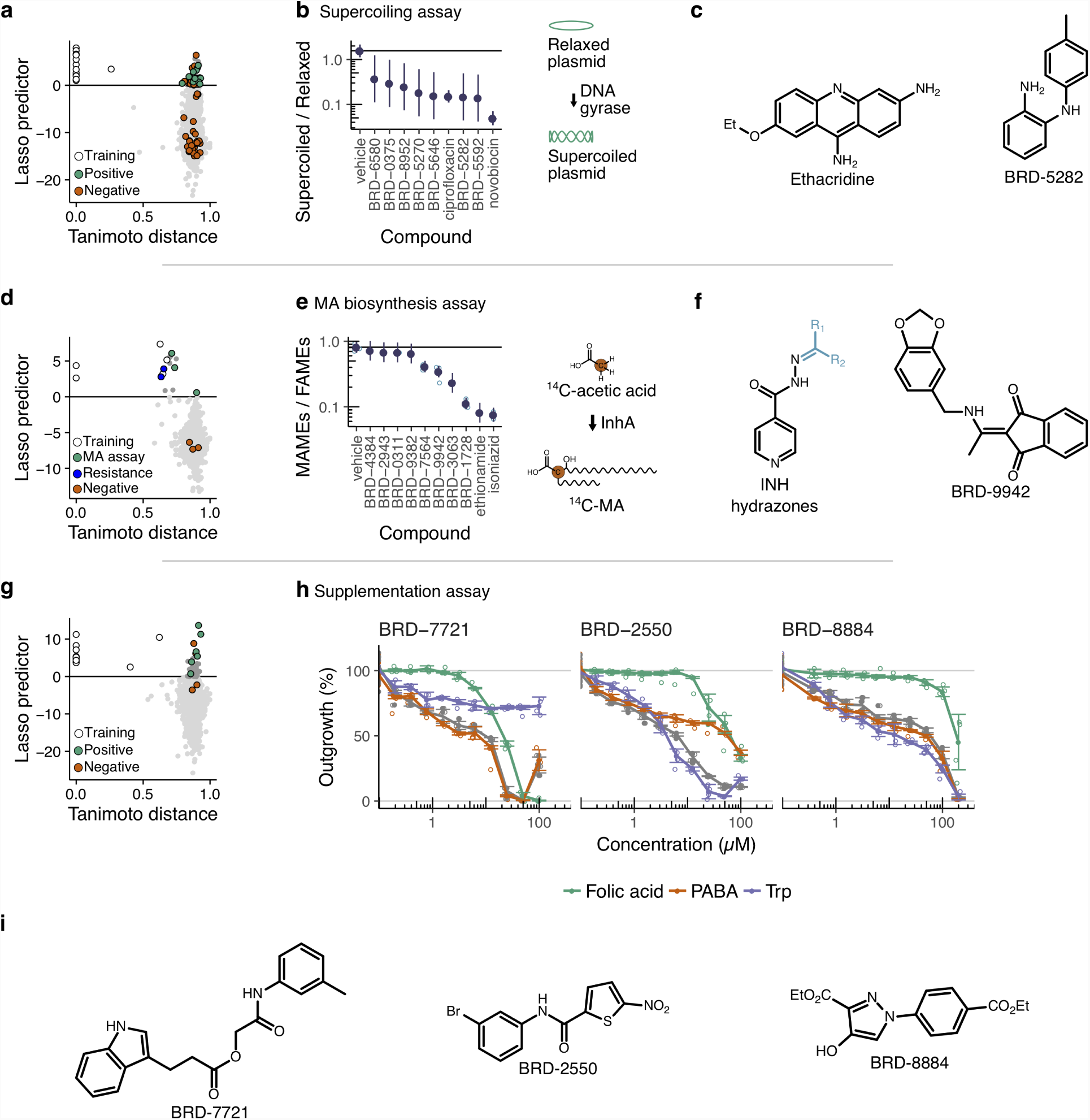
New inhibitor classes of well-validated targets in M. tuberculosis. **a**, Scatter plot of Lasso predictor (strength of prediction) for DNA gyrase inhibition against Tanimoto distance (chemical structure dissimilarity) to the fluoroquinolone family of known DNA gyrase inhibitors. Each point is a compound in the screen. All compounds above the horizontal line were predicted to be DNA gyrase inhibitors. White points were known fluoroquinolone DNA gyrase inhibitors in the training set, with compounds confirmed in an in vitro assay for DNA gyrase inhibitory activity are shown in green. Compounds with no in vitro activity are shown in orange. **b**, Actual compound performance of predicted DNA gyrase inhibitors in an in vitro assay of DNA gyrase supercoiling inhibition. We tested for inhibition of DNA gyrase supercoiling activity in an agarose gel-based assay. The ratio of imaged pixel intensities for supercoiled and relaxed bands was indicative of inhibition activity, as shown by the ciprofloxacin positive control. Compounds that showed statistically significant (*p* < 0.05, Wald test; *n* = 2) inhibition are shown. Error bars show standard errors of the GLM regression coefficients. **c**, Examples of new DNA gyrase inhibitor chemotypes predicted by the Lasso classifier and confirmed in vitro. **d**, As (a), but for the mycolic acid biosynthesis classifier. White points were known mycolic acid biosynthesis inhibitors in the training set, with green points indicating compounds confirmed to inhibit 14C-acetic acid incorporation into mycolic acid; Mtb with a loss-of-function mutation in KatG (KatG^−^) or InhA expression (BAA-812) were resistant to compounds shown in blue. **e**, Actual compound performance of predicted mycolic acid biosynthesis inhibitors in an in vitro assay of inhibition of 14C-acetic acid incorporation into mycolic acid. The ratio of imaged pixel intensities for fatty acid methyl esters (FAMEs) and mycolic acid methyl esters (MAMEs) bands was indicative of inhibition activity, as shown by the isoniazid and ethionamide positive controls. Error bars show the 95% confidence interval (*n* = 2) of the GLM regression coefficients. **f**, New mycolic acid biosynthesis inhibitor chemotypes predicted by the Lasso classifier and confirmed *in vitro*. **g**, As (a), but for the folate classifier. White points were known sulfonamide folate biosynthesis inhibitors in the training set, with green points indicating compounds whose growth inhibitory activity was abolished by PABA or folic acid supplementation. Compounds whose growth inhibitory activity was not abolished by PABA or folic acid supplementation are shown in orange. **h**, Actual compound performance of predicted folate biosynthesis inhibitors in a metabolite rescue assay. Mtb was treated with predicted inhibitors in the presence or absence of tryptophan, folate, or PABA. The effect of BRD-7721, a 3-indole propionic acid ester, is abolished by tryptophan supplementation, indicating it is a tryptophan biosynthesis inhibitor. In contrast, the effect of BRD-2550 (a nitrothiophene) is abolished by folate and PABA, while that of BRD-8884 is abolished by folate alone, showing that they are inhibitors of folate biosynthesis with distinct mechanisms. Individual replicates (*n* = 4) are shown as open circles, means are shown as filled circles, and error bars show 95% confidence intervals. **i**, Examples of new folate and tryptophan biosynthesis inhibitor chemotypes predicted by the Lasso classifier and confirmed by metabolite supplementation.

Using an Mtb DNA gyrase supercoiling and decatenation in vitro assay, we confirmed 27 (52%) of 52 predicted new DNA gyrase inhibitors (Fig. 3b). In contrast, 25 randomly selected compounds showed no activity, showing that the classifier significantly enriched for real inhibitors (*p* = 2 × 10^−7^, Fisher’s exact test; ROC AUC = 0.89). Of the validated compounds, ethacridine’s acridine scaffold (Fig. 3c) had been previously reported to inhibit Mtb DNA gyrase (29). The model also predicted tryptanthrin, an antiinfective whose target has eluded extensive antibacterial and antitrypanosomal research (30, 31); we validated it to be a DNA gyrase inhibitor (Extended Data Fig. 4). All remaining scaffolds were novel.

### Mycolic acid biosynthesis inhibitors

The active forms of the cornerstone clinical antitubercular prodrugs INH and ETH both inhibit InhA (32), a key enzyme in mycolic acid biosynthesis. We sought new inhibitors of this pathway by training on chemical-genetic interaction profiles of these drugs. Although the strain pool did not include an InhA hypomorph, our analysis nevertheless yielded an excellent predictive model (Fig. 3d) using increased relative fitness of the MshC sentinel strain as the most discriminatory feature (Extended Data Fig. 3b). MshC catalyzes the incorporation of cysteine into mycothiol, an antioxidant unrelated to mycolic acid biosynthesis, whose depletion has been shown to confer resistance to INH and ETH (33).

Further validating our approach, the model predicted six hydrazone derivatives of INH. A clinical loss-of function mutant of catalase KatG, which activates the INH prodrug, was resistant to the six INH-hydrazones, indicating that the INH-hydrazones are activated in the same way as INH. Importantly, the model also predicted one completely novel scaffold, the indenedione BRD-9942. Since these compounds had measurable activity against wild-type Mtb, we found that, like INH and ETH, the INH-hydrazone scaffold and BRD-9942 inhibited 14C-acetate incorporation into mycolic acids (Fig. 3e-f).

### Folate and tryptophan biosynthesis inhibitors

Because inhibiting folate biosynthesis is an effective antimicrobial strategy against many bacteria, but not yet exploited against Mtb, we sought new classes of inhibitors against this pathway. Training on the chemical-genetic interaction profiles of the sulfonamides, which are known dihydropteroate synthase (DHPS) inhibitors, the most discriminatory feature was inhibition of the TrpG hypomorph (Extended Data Fig. 3c). TrpG is involved in both folate and tryptophan biosynthesis (Extended Data Fig. 5a), catalyzing formation of both 4-amino-4-deoxychorismate (a folate precursor) and 2-amino-2-deoxyisochorismate (a tryptophan precursor).

We tested whether the inhibitory effects of 7 of the 43 predicted compounds (Fig. 3g), spanning several chemotypes and with measurable inhibition of wild-type Mtb, could be abolished by supplementation with tryptophan, folate, or the folate pathway intermediate para-amino benzoic acid (PABA) (Fig. 3h). The nitrothiophene amide or ester compounds, BRD-2550, BRD-3387, BRD-5592, and BRD-9737 (Fig. 3i; Extended Data Fig. 5e), as well as a derivative of para aminosalicylic acid (PAS), BRD-9819, behaved similarly to methotrexate and PAS (known DHFR inhibitors; Extended Data Fig. 4b), with their effects abolished by both PABA and folate supplementation. In contrast, BRD-8884 had effects that could only be rescued by folate and not PABA (Fig. 3h-i), suggesting inhibition of a novel, late step in the folate pathway. Thus, this strategy identified inhibitors targeting different enzymatic steps in the folate biosynthetic pathway.

Finally, BRD-7721, a 3-indolepropionic acid (3-IPA) ester predicted to be a folate biosynthesis inhibitor, was only rescued by tryptophan supplementation (Fig. 3i-h), indicating that it inhibits tryptophan biosynthesis. 3-IPA was recently identified as having antimycobacterial activity in a fragment-based screen (34); the activity of this free acid was also abolished by tryptophan supplementation.

Based on our empirical validation of new folate and tryptophan inhibitors, we trained two new models: an updated folate biosynthesis inhibitor predictor and a tryptophan biosynthesis inhibitor predictor. The refined classifier weights the behavior of the FolB hypomorph in conjunction with TrpG in its discrimination of folate inhibitors (Extended Data Fig. 3d-e).

## Screening a larger, unbiased compound library

We applied PROSPECT to a large unbiased library of ∼50,000 compounds at 50 µM and a 152-strain pool (including 94 of the 100 strains used previously), to demonstrate its scalability, to identify compounds whose potential activity against wild-type Mtb would be revealed by hypersensitive hypomorphs, and to demonstrate the ability to identify inhibitors of novel targets with potent activity against wild-type Mtb after medicinal chemical optimization. Of the7,245,009 chemical-genetic interactions tested, 95,685 (1.3%) were strongly inhibitory. Selecting 1331 compounds for retest-ing, we confirmed 78% of the inhibitory chemical-genetic interactions (LFC < 0), resulting in an ROC AUC of 0.74, similar to that observed for the first, smaller screen. In an orthogonal growth assay, the ROC AUC was 0.69 (Fig. 4a).

**Figure 4:**
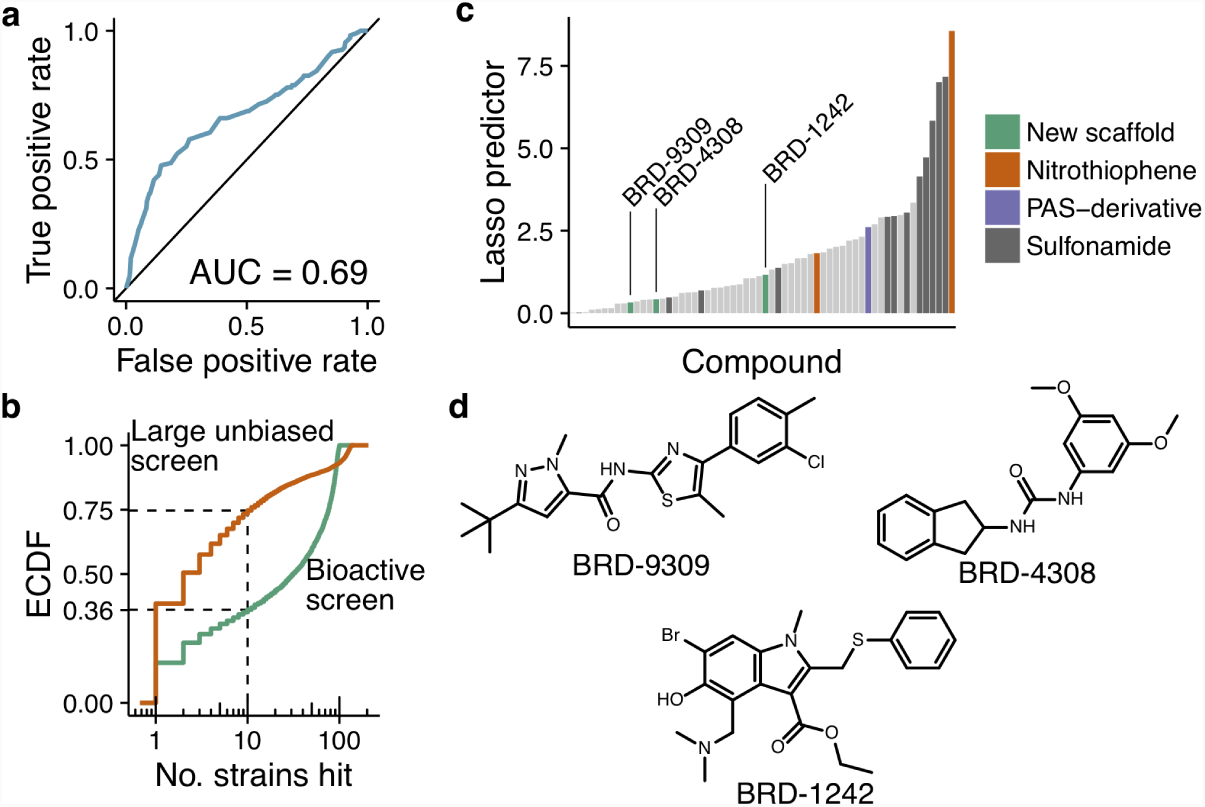
Performance of a larger, 50,000-compound screen. **a**, ROC curve of primary data against a confirmatory secondary assay. We retested more than 100 compounds predicted to have activity in the primary screen using a resazurin, growth-based colorimetric assay. Taking 75% inhibition in the secondary assay as ground truth, we demonstrated the primary assay as predictive of real activity that could be detected by conventional methods. **b**, Compounds in the library of bioactive compounds generally hit more strains than compounds in the unbiased library. Empirical cumulative distribution functions of number of hypomorphs hit by compounds in the two screens is plotted. Shown by the dotted lines, 36% of compounds in the bioactive library and 75% of compounds in the larger library hit 10 strains or fewer, suggesting that activity detected in the larger screen was generally more hypomorph-specific. **c**, Lasso predictor scores from the folate biosynthesis inhibitor classifier applied to the large unbiased screening data. The highest-scoring compounds were known folate inhibitors (sulfonamides and nitrothiophene compounds), thus validating the approach. New scaffolds were also identified. **d**, Three new folate biosynthesis inhibitor chemotypes predicted by the Lasso classifier and confirmed by metabolite supplementation.

The hit rate against wild-type Mtb alone was 0.9% (436 compounds), typical for an unbiased whole-cell compound screen; in contrast, 10-fold more compounds (4403; 9%) were active against at least one of the 152 hypomorph strains (Extended Data Fig. 6a). Additionally, 92 of the 152 hypomorphs had the most negative LFC for at least three compounds, suggesting high MOA diversity in this unbiased library. Of the 4403 active compounds, 3967 had no activity against wildtype Mtb. However, 73% were highly specific against the hypomorphic strains (1-10 strains hit) and 11% were moderately specific (11-50 strains hit); the remaining 16% were relatively non-specific (>50 strains hit). This distribution shows greater specificity than for the compounds in the first, bioactive library (35%, 31%, and 34%), likely because smaller library is enriched for compounds with wild-type Mtb activity (Fig. 4b). The larger library also yields a greater diversity of chemical-genetic interaction profiles, suggesting greater target diversity, as evidenced by hierarchical clustering of the profiles (1864 distinct clusters in the unbiased library vs. 235 in the bioactive library) (35). That the profile clusters in the unbiased library are meaningful is supported by the fact that over one-third of them are enriched for structurally similar compounds (Extended Data Fig. 6b-c).

We applied the folate biosynthesis inhibitor classifier derived from training on the smaller, bioactive library to the data from the unbiased library. Despite being trained on a dataset generated from multiple concentrations of each compound and a smaller hypomorph set, the folate model showed excellent transferability, predicting 60 compounds from this larger screen (Fig. 4c), including twelve sulfonamide analogues, one derivative of PAS, and two nitrothiophene amides, which we have shown to inhibit the folate pathway (Fig. 3i). Three compounds (Fig. 4d) represent additional novel scaffolds that we validated as also acting in the folate pathway based on suppression of their activity with the addition of PABA or folate (Extended Data Fig. 6d). These results thus demonstrate the scalability and generalizability of this strategy, and the potential to leverage the much higher hit rate and wider target space obtained by performing primary screening with a hypomorph library to provide new chemical scaffolds against novel targets.

## Discovering inhibitors without reference data

In the examples above, we were able to rapidly identify new chemotypes against established targets based on reference data derived from compounds with known MOAs. However, discovering first-in-class molecules with completely novel MOAs and targets requires identifying inhibitors without available reference data. We therefore developed an approach to identifying inhibitors of a target protein of interest based on specific inhibition of the corresponding hypomorphic strain.

As an initial test, we applied it to the discovery of new inhibitors of RNA polymerase (RNAP) — a high priority target of the rifamycins, which anchors antitubercular regimens but for which there is rising resistance. While RNAP inhibitors exist, our screen provided no information about their chemical-genetic profiles because the concentrations of rifamycin in our screen inhibited growth of all strains. Instead, we looked for compounds that showed significant specificity for the RpoB hypomorph, which is depleted for RNAP.

We prioritized 20 compounds with strong chemical-genetic interaction with the RpoB hypomorph (*p* < 10^−10^), requiring it to be among the two most inhibited strains for at least one dose. Testing these compounds in an in vitro RNA polymerase assay (36, 37), three compounds – including the antineoplastic human RNAP inhibitor actinomycin D – showed direct inhibition of E. coli RNAP (Fig. 5a-b). While the positive predictive value was lower than using machine learning, the approach readily identified new scaffolds against this important target, which has proved recalcitrant to the discovery of new inhibitors through whole cell screening (38).

**Figure 5:**
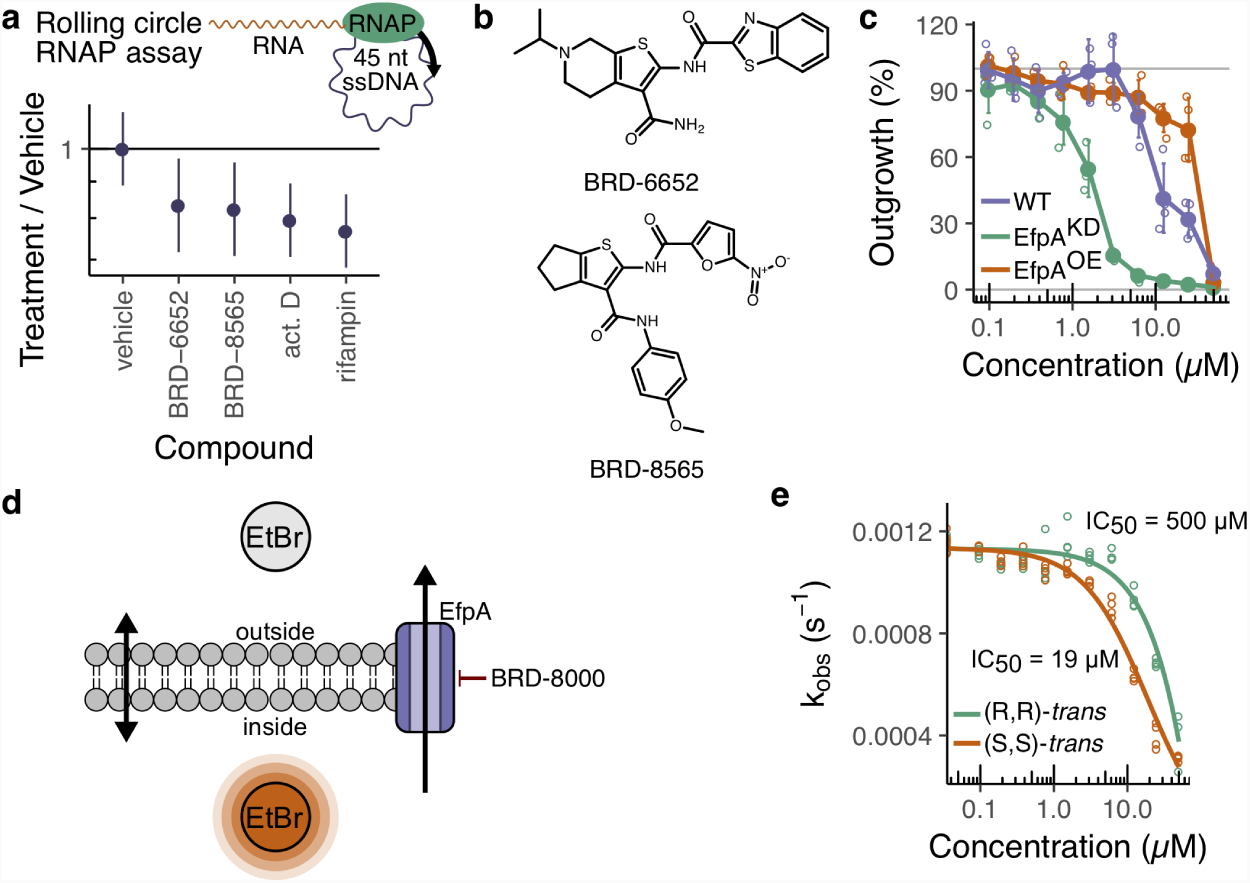
Discovery of new inhibitors in the absence of reference data, including inhibitors of a novel target in *M. tuberculosis.* **a**, Actual compound performance of predicted RpoB inhibitors in an in vitro assay for inhibition of RNA synthesis by *E. coli* RNAP. Three compounds which showed statistically significant inhibition are shown with a rifampin control (*p* < 0.05, two-tailed Wald test; *n* = 4). Error bars show the 95% confidence interval of the GLM regression coefficients. Act. D: actinomycin D. **b**, New RNAP inhibitor chemotypes validated in vitro. **c**, Dose response of BRD-8000 on growth of wild type Mtb, the EfpA hypomorph, and a mutant overexpressing EfpA, demonstrating hypersensitivity of the hypomorph. Individual replicates (*n* = 4) are shown as open circles, means are shown as filled circles, and error bars show 95% confidence intervals. **d**, Schematic of the EtBr efflux assay. Bacteria were loaded with EtBr and its efflux was monitored by change in fluorescence. **e**, Measurements of first-order rate constant for EtBr efflux, *k*_*obs*_, against varying concentration of BRD-8000.2 enantiomers. Inhibition of EtBr efflux by BRD-8000.2 is enantiospecific, with level of efflux inhibition correlating with Mtb growth inhibition. Individual replicates are shown as open circles, and fitted curves are shown as lines.

## Discovering inhibitors of a novel target

Finally, we demonstrated that PROSPECT can identify inhibitors against a completely novel target that would not be found by conventional strategies. Specifically, we identified a compound that inhibits a new target, EfpA and optimized it to achieve potent activity against wild-type Mtb.

We analyzed the first screen for compounds that did not strongly inhibit wild-type Mtb, but were strongly active against at least one hypomorph at all screening concentrations. These were then ranked by how few hypomorphs were significantly inhibited – a proxy for specificity – and whether the compound was a chemically attractive scaffold. The highest ranked interaction was BRD-8000 with the hypomorph of EfpA, an uncharacterized essential efflux pump. Measurement of the compound’s MIC_90_ confirmed that BRD-8000 has strong activity against the EfpA hypomorph (MIC_90_ = 6 µM) with little wild-type activity (MIC_90_ ≥ 50 µM) (Fig. 5c).

We optimized BRD-8000, first by resolving the mixture of stereoisomers present in the initial chemical library; the (S,S) trans stereoisomer is the active isomer (BRD-8000.1, egy, and the potential to leverage the much higher hit rate and wider target space obtained by performing primary screening Table 1), with a wild-type MIC_90_ of 12.5 µM. Migrating with a hypomorph library to provide new chemical scaffolds against novel targets. the pyridyl bromine from the 6-to the 5-position to obtain BRD-8000.2 maintained hypomorph hypersensitivity while improving wild-type MIC_90_ potency (MIC_90_ = 3 µM, Table 1). We generated more than 30 independent resistant mutants against BRD-8000.2 in wild-type Mtb (resistance frequency of ∼10^−8^); all mutants contained the same C955A mutation in efpA (EfpA_V319F_), thereby providing genetic support for EfpA as the target. Further chemical optimization yielded BRD-8000.3, a methyl-pyrazole derivative of BRD-8000.1 with MIC_90_ of 800 nM against wild-type Mtb, a ≥ 60-fold overall improvement in activity from the original hit (Table 1).

**Table 1:**
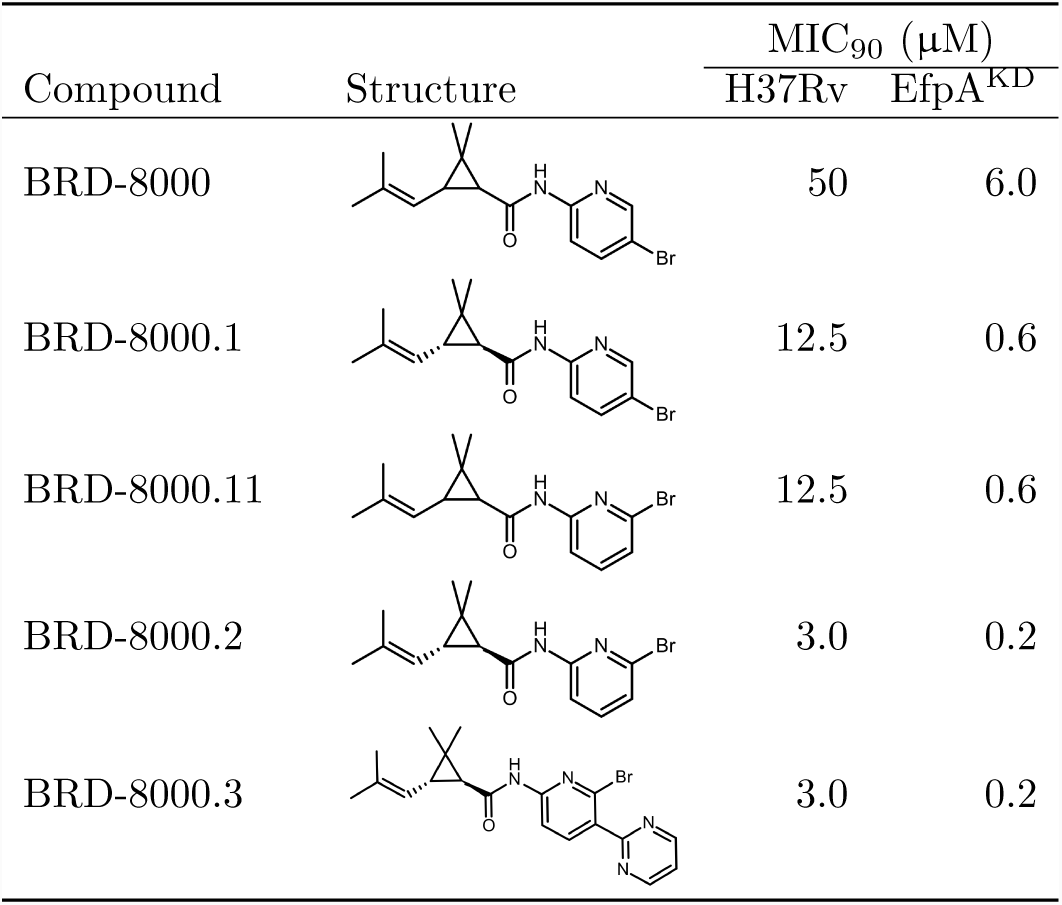
Potencies of compounds targeting the essential efflux pump EfpA. MIC_90_: Minimum inhibitory concentration at 90%; EfpA^KD^: EfpA hypomorph.

| Compound | Structure | MIC<sub>90</sub> (μM) |  |
| --- | --- | --- | --- |
|  |  | H37Rv | EfpA<sup>KD</sup> |
| BRD-8000 |  | 50 | 6.0 |
| BRD-8000.1 |  | 12.5 | 0.6 |
| BRD-8000.11 |  | 12.5 | 0.6 |
| BRD-8000.2 |  | 3.0 | 0.2 |
| BRD-8000.3 |  | 3.0 | 0.2 |

To functionally confirm that the BRD-8000 series targets EfpA, we took advantage of the fact that BRD-8000 does not kill *Mycobacterium smegmatis* (Msm), likely because EfpA is not essential in Msm(39), to investigate the series’ effect on efflux through EfpA (Fig. 5d). BRD-8000 and BRD-8000.2 inhibited efflux of ethidium bromide (EtBr), a known substrate of EfpA (39), with an IC50 of 38 µM and 15 µM, respectively, reflective of their corresponding MIC_90_ values (Extended Data Fig. 7a). This inhibition was stereospecific as the inactive (R,R)-trans isomer of BRD-8000.2 has an efflux IC50 of 500 µM (Fig. 5e). Finally, BRD-8000.2 inhibits EfpA by an uncompetitive or non-competitive inhibitory mechanism as inhibition is independent of EtBr concentration (Extended Data Fig. 7b), and has a high-affinity interaction with EfpA that is eliminated from Msm when efpA is deleted. This highaffinity interaction is restored by episomal complementation of Mtb’s efpA homolog (Extended Data Fig. 7c).

The BRD-8000 series is bactericidal (Extended Data Fig. 7d), kills non-replicating phenotypically drug-tolerant Mtb (MBC50 = 390 nM), has low human toxicity (hepatocyte IC50 = 100 µM), and is narrow-spectrum, not inhibiting growth of *E. coli, S. aureus, Pseudomonas aeruginosa*, or *Klebsiella pneumoniae*, while the MIC_90_ for *M. marinum* was 25 µM. Having confirmed that BRD-8000 indeed inhibits the novel target EfpA, we returned to the primary screening data to identify additional EfpA inhibitors based on chemical-genetic interaction profile similarity. We tested 11 prioritized compounds for their ability to inhibit EtBr efflux and identified three new scaffolds, encompassing six molecules, that inhibited EtBr efflux (Extended Data Fig. 7e). While not as specific for the EfpA efflux pump as BRD-8000, given that EfpA is the only essential efflux pump in Mtb (17), these compounds’ whole cell activity is likely due to their activity on EfpA. In particular, BRD-9327 does not act competitively. Thus, by taking an iterative approach, we were able to use PROSPECT to quickly expand the chemical diversity of small molecule candidates against a novel target.

## Discussion

We have developed PROSPECT, a powerful and rapid chemical-genetic interaction profiling strategy, which is able both to discover many new potential compounds for Mtb drug development and to gain insight into their MOA from the primary screening data. By allowing immediate integration of potential target information into hit prioritization, it shifts hit selection away from the conventional approach of relying simply on compound potency. Interpreting this complex, multidimensional data guided either by the chemical-genetic interaction profiles of known compounds or, in the absence of reference data, a manner that relies on hypomorph specificity, we rapidly identified MOA for 45 new molecules. Importantly, these hits included new scaffolds against known targets – a valuable strategy to overcome antimicrobial resistance (40), and the first inhibitors against completely new targets, taking advantage of the enormous but underexplored target space in Mtb as revealed by genomic studies. With its ability to identify ∼10-fold more hits than obtained by screening wild-type Mtb alone, PROSPECT greatly expands both the chemical and target space of identified compounds.

We demonstrate how PROSPECT can identify Mtb small molecule candidates against a novel target that could not have been discovered by conventional approaches with the discovery of an EfpA with potent wild-type activity after chemical optimization of the initial hit compound, a process similar to the optimization required in conventional approaches to improve the potency and drug like characteristics of initial hit compounds. Importantly, mining of the entire chemical-genetic interaction dataset can be performed iteratively, with the discovery of inhibitors of new targets, such as BRD-8000, enabling the discovery of additional inhibitors (BRD-9327) against the same target by chemical-genetic interaction profiles for scaffold hopping. While only the surface has been scratched thus far for these large datasets, we have provided examples how integrating MOA insight into primary whole-cell screening can transform the targets and molecules that emerge and are prioritized. With this in mind, we have made primary data publicly available (broad.io/cgtb) to catalyze the entire community’s discovery of new inhibitor classes and their respective targets and propose that PROSPECT is widely applicable to important pathogens beyond Mtb.

## Acknowledgements

Funding was provided by Bill and Melinda Gates Foundation, Broad Institute TB Gift Donors and Pershing Square Foundation.

## Author contributions

The manuscript was written by E.O.J., E.S.L., and D.T.H. Statistical analysis was carried out by E.O.J. Computational pipelines were written by E.O.J. and N.B. Experiments were designed as follows. Strain construction: K.M.G., K.C.M., T.R.I., S.E., E.J.R., C.M.S, D.S. Assay development and screening: E.O.J., J.E.G., M.F., D.T.H. Medicinal chemistry: T.K., B.K.H., M.H.S.-W., D.T.H. Mechanism of action follow-up: E.O.J., J.E.G., D.T.H. Experiments were carried out as follows. Strain construction: R.E.A., N.B., I.D.S., M.G., J.A.M., C.E.M., K.G.P., J.T.P., P.A.P., M.K.P., N.R., N.S., C.T., S.W., J.B.W. Assay development: E.O.J., M.T. Compound screening: E.O.J., E.L., M.S., K.D., J.D., C.G., A.G., R.K., R.N., M.T., C.W. Mechanism of action follow-up: E.O.J., J.E.G., E.O., E.M.

