## Supplementary material for "Large-scale chemical-genetics yields new *Mycobacterium tuberculosis* inhibitor classes": Methods and supplementary figures

**Supplementary Materials:**

Methods

Extended Data Figures 1-7

### Methods

#### Strain selection and construction

To create strains of Mtb with reduced levels of target proteins, we employed a protein degradation system previously described[1]. Briefly, a DAS+4 tag (abbreviated as DAS-tag) was recombineered into the chromosome of Mtb H37Rv, at the 3′-end of the target gene. Next, the DAS-tagged mutant was transformed with a plasmid containing *sspB* downstream of an inducible promoter. When induced, SspB delivers DAS-tagged protein to the native caseinolytic protease ClpXP for degradation. In order to generate hypomorph strains with varying levels of knockdown, we developed plasmids producing graded levels of SspB (Extended Data Fig. 1a). This was achieved by varying the strengths of both the promoter driving transcription of *sspB* and the translational initiation signal required to produce SspB protein. Regulation was achieved by repression of the *sspB* promoter by a reverse tetracycline repressor (revTetR). RevTetR requires anhydrotetracycline (ATC), which acts as a corepressor, to shut down transcription of *sspB*. Repression of *sspB* suppresses degradation of the DAS-tagged protein. Phenotypically we thus refer to these mutants as TetON mutants (because the presence of ATC represses degradation of the DAS-tagged target protein).

To facilitate the large scale of our approach, a sequence-design program was developed (http://orca2.tamu.edu/tom/U19/seqtool.html), which assembled the sequences of recombineering cassettes automatically. Every cassette consists of 500 bp flanking sequences around the stop codon of the target, the DAS tag (inserted at the 3′-end of the target gene), a loxP site, a unique nucleotide sequence ("molecular barcode"), and a hygR selectable marker. If the target gene was located less than 21 bp upstream of the 5′-end of an adjacent ORF then a new ribosomal binding site was inserted to preserve translation of the downstream gene. The designed DNA fragment was synthesized (Gen9, Cambridge, MA, or GenScript, Piscataway, NJ) in plasmid pUC57 with flanking PmeI sites. The fragment was excised from the plasmid with PmeI and used as a double-stranded DNA recombineering substrate [2].

Molecular barcodes enabled identification and quantification of each strain amongst a pool of strains. Each barcode region was 74 nucleotides long, with common flanking regions on each end that include a PacI site (underlined) and primers for PCR amplification (italics), and a unique sequence of 20 nucleotides in the middle (<20N>), which is the barcode: ttaattaATCTTGTGGAAAGGACGA<20N>ACGCTATGTGGATACGCTGCTTTAattaa. Each barcode is unique to each target, thus only one SspB version strain of any target can be included in a given pool.

#### Compound library assembly

The library was drawn from a pool of 1128 compounds we had identified in internal screening efforts as having activity against wild-type Mtb H37Rv[3-5], 5611 compounds reported by Southern Research Institute[6-8], and 177 compounds reported by GlaxoSmithKline[9]. After removing duplicates, we assembled a library of 2000 available compounds. We supplemented this collection with 1226 compounds from Selleck's Pharmacologically Active Compound Library, to include known antibiotics to serve as positive controls and as ground truth data for machine learning. To confirm the reported Mtb activity, we screened the library against GFP-expressing wild-type Mtb H37Rv and found that 1312 (45%) had an $MIC_{90}$ < 64 μM.

This library was assembled from internal Broad Institute chemical libraries, including sub-libraries from the Broad diversity-oriented synthesis collection [10], National Institutes of Health's Molecular Libraries Probe Production Centers Network library, and purchased from WuXi, filtered for physico-chemical properties including molecular weight < 550, calculated log(D) at pH 7.4 < 4, polar surface area < 110, charge at pH 7.4 > –1.

**Optimization of screening conditions and analysis**
We optimized screening assay parameters in 384-well format, including compound exposure time (14 days), genomic DNA extraction (10% dimethylsulfoxide (DMSO) and 95ºC for 15 min), and PCR conditions (20 cycles, annealing at 65 ºC) that maximized the robust $Z'$-score of log-transformed counts for a RIF dilution series and minimized random noise as determined by strain-wise coefficient of variance (CV).

An appropriate sequencing depth (500 reads per strain per well) was determined by performing repeated simulations of a 10,000-well HTS dataset with active compounds present at 1%, balancing cost, accuracy, sensitivity, and inference of fitness via a generalized linear model (GLM).

**Multiplexed screening of compound libraries**
Strains of the final screening pool were grown separately in Middlebrook 7H9 (Difco) supplemented with oleic albumin dextrose catalase (OADC, from Becton Dickinson) and 10 mM sodium acetate, appropriate antibiotics, and 1 μg/mL ATC. When the cultures reached mid-exponential growth phase, $OD_{600}$ was measured and bacteria were combined equally into a single pooled culture, which was then diluted in Middlebrook 7H9-OADC-acetate to an approximate $OD_{600}$ of 0.005. This culture was washed three times in Middlebrook 7H9-OADC-acetate to remove ATC. 40 μL dilute culture was distributed into wells of clear polystyrene 384-well plates (Corning), which contained 1 nL of screening compound per well as prepared by Broad Institute Compound Management. On every plate, rows A, B, O, and P and columns 1 and 24 were left empty to prevent edge effects arising from evaporation. Columns 2 and 23 were occupied by alternating DMSO (negative vehicle) and rifampin on-board controls. Each batch of screening also included eight control plates (four inoculated at the beginning of the day and four at the end) which contained 12-point two-fold serial dilutions of rifampin and trimethoprim, and for the larger screen BRD-4592 and methotrexate in addition.

Plates were incubated for 14 days in humidified containers at 37 ºC. 40 μL 10 % v/v aqueous DMSO was then added to each well, before the plates were decontaminated by heating at 80 ºC for 2 h. One PCR was performed per well in 384-well PCR plates (Eppendorf) containing 1 μL heat-inactivated culture, 5μL 2× Q5 Master Mix (NEB), 0.25 μL forward and reverse primers, 1 μL 10x Q5 buffer, and 2.5 μL MilliQ water. The primers contained 5′-overhangs which added plate and well identification barcodes as well as nucleotide sequences necessary for Illumina NGS (Extended Data Fig. 1c). PCR was carried out as recommended by NEB for 20 cycles, using a 2 min extension time and 65 ºC annealing temperature.

5µL samples from each PCR were combined into a single pool; unused primers were twice removed using AMPure XP beads (Beckmann) at 2× the pooled PCR volume, finally eluting in 200 µL MilliQ water. Final sequencing library quality control was carried out using a Bioanalyzer High-Sensitivity DNA kit (Agilent).

Sequencing was carried out at the Broad Genomics Platform using Illumina HiSeq 2500 at a sequencing depth of at least 500 reads per strain per well.

**Barcode counts from Illumina NGS**

The ConCensusMap script was written in Python. Since sequencing reads had a consistent structure, the script, provided with the locations of barcodes within each sequencing read, takes as input the undemultiplexed main FASTQ file and index FASTQ file and counts the co-occurrence of each combination of the three barcodes corresponding to plate, well, and strain. These counts are then annotated with compound information based on the inferred plate and well coordinates and strain identity based on the strain barcode. The output is a comma-separated value (CSV) file with one line per strain and well combination.

**Fitness inference from barcode counts**

In order to determine an effective analysis method and the depth of sequencing required, we first noted that counts from DMSO treated control wells appeared to be drawn from a negative binomial (NB) family distribution. We then repeatedly simulated ideal HTS datasets by drawing from a pseudo-random NB distribution. We chose to perform these simulations under conditions reflecting a typical low prevalence compound screening scenario, setting hit compounds with 50% inhibition activity being present at one percent, in anticipation of screening large, unbiased libraries. We did so with the understanding that the analysis method and depth of sequencing that would be required in the more general, stringent case would more than suffice in the case of the specific library enriched for TB active compounds. We found that a NB family generalized linear model (GLM) provided the most consistent specificity and sensitivity at lower sequencing depths and suggested a sequencing depth of 500 reads per strain per well to be an ideal balance of cost, accuracy, and sensitivity. Conveniently, the GLM framework also allowed dynamic correction of systematic variation (or batch effects) of sequencing data; the strain-wise NB dispersion nuisance parameter was estimated by maximizing profile likelihood as described previously [11]. The analysis protocol and quality control checks were developed into a pipeline called ConCensusGLM, which generates an estimated $\log_2$(fold change) (LFC) of counts compared to the DMSO control screening wells, and an estimated two-tailed *p*-value (Wald test) for each unique strain, compound, and compound concentration combination. This LFC value is directly related to change in fitness (doubling time) of a strain on exposure to a compound and reflects the chemical genetic interaction between a compound and strain.

Estimation of changes in counts between control and test conditions has been implemented before, notably for RNA-seq[11,12]. The present task was similar but was complicated by two problems.

Firstly, the assumption of previous implementations for RNA-seq was that the abundance of most transcripts does not change between conditions; this assumption is used for normalization. Here, high concentrations of an active compound will cause all strain abundances to be close to their inoculum, i.e. very different from an untreated DMSO reference. Applying such an assumption to our data would result in throwing out information on very potent compounds.

Secondly, the number of test conditions could be potentially very large and would be spread across batches. We therefore sought a principled way to model batch-to-batch variation in a computationally efficient manner.

The script ConCensusGLM tackles these issues simultaneously. Since observed counts $K_{sci}$ of a strain s in well $i$ and condition $c$ could be modeled as a Negative Binomial (NB) distributed random variable (i.e. $K_{sci} \sim \mathrm{NB}(\mu_{sci}, \alpha_s)$ where $\mu_{sci}$ is the true unobserved mean count for well $i$ of strain $s$ in the presence of condition $c$, and $\alpha_s$ is the strain-wise dispersion parameter), we turned to a NB family generalized linear model (GLM) with a log link to estimate log fold change of counts in a given condition compared to an untreated reference (*58*). It was natural for us, therefore, with the advantage of many negative control data points, to include experimental metadata as additional GLM predictors and isolate the effect of the compounds: $\log \mu_{sci} = \beta_{0s} + x_{ci}\beta_{sc} + y_{sq(i)}\omega_{sq(i)}$ where $\beta_{0s}$ is the regression intercept (interpreted as the mean count for strain $s$ in DMSO negative control wells), $x_{ci}$ is the indicator variable indicating presence and absence of condition $c$ in well $i$, $\beta_{sc}$ is the regression coefficient of condition c (interpreted as log fold change), and $y_{sq(i)}\omega_{q(i)}$ is the product of the indicator variable indicating experimental metadata $q(i)$ and its regression coefficient.

However, with many conditions, the GLM design matrix became computationally unwieldly. To address this, a minimal negative binomial GLM per strain (i.e. $\log \mu_{s0i} = \beta_{0s} + y_i\omega_{q(i)}$) was fitted using iteratively weighted least squares (IWLS) to only the DMSO negative control wells, using recorded experimental metadata, such as experiment date, which thermocycler, which sequencing lane, and which plate as categorical predictors. Rough method of moments estimates of the strain-wise dispersion nuisance parameters $\alpha_s$ were used in the model. The models were then used to estimate the expected counts $\gamma_{sq(i)} = \exp(\beta_{0s} + y_{si}\omega_{sq(i)})$ (termed offsets) for any given strain in any given well $i$ in the screen based on that well's set of metadata $q(i)$.

Secondly, the final strain-wise dispersion parameters $\alpha_s$ were estimated by maximizing Cox-Reid adjusted profile likelihood across the whole dataset following the procedure in [11] but including the offsets when calculating likelihood. This alteration prevented loss of power arising from positive bias in dispersion estimates when batch effects are not accounted for. Simulations indicated that this approach produced consistent and unbiased estimates of dispersion.

Finally, one GLM was fitted per compound (to minimize the dimensions of the design matrix and afford parallelization) per strain (using the final strain-wise dispersion estimates) without an intercept but using the offsets (i.e. $\log \mu_{sci} = x_{ci}\beta_{sc} + \log \gamma_{sq(i)}$). Quoted log fold changes (LFC) and $p$-values are from these final models as implemented in R[13] with the MASS package[14].

**Unsupervised machine learning**

The compound target ground truth dataset was identified through a combination of extracting annotations from the KEGGDrug database[15] and manual curation. The 400-dimensional data were visualized using *t*-SNE[16], as implemented in the Rtsne package[17] for R.

Hierarchical clustering of chemical-genetic interaction profiles was performed using the Ward's algorithm[18] as implemented in R's hclust function. The number of clusters in the data was estimated using the Gap statistic[19].

Enrichment of chemical structure similarity within a given chemical-genetic cluster was calculated as the difference between the within-cluster sum-of-squares of the Tanimoto distances

(Tanimoto WSS), and the mean Tanimoto WSS ($n$ = 10,000) for sets of the same number of randomly chosen compounds from the same compound library.

**Supervised machine learning**

Despite the variety of putative targets represented in the ground truth compound set, only a few mechanisms were well characterized and had enough representatives to be used for training and validation. In addition, some classes, such as the penicillins, despite being well characterized and represented, were considered inappropriate classifier labels since they have no activity in Mtb due to intrinsic inactivation pathways. However, we wanted to be able to incorporate the negative information that such compounds carried; for example, it is reasonable to assume that ampicillin is not a protein synthesis inhibitor. To take this into account, we constructed a binary classifier for each mechanism of interest: known inhibitors from the ground truth set of a target of interest were labeled as "true" and the rest of the ground truth set was labeled "false".

Models were implemented as Lasso-regularized binomial-family (with logistic link function) GLMs using the glmnet package [20] for R. For model training, 40% of the ground truth compounds' data was held out for validation. The remaining 60% was subjected to cross-validation to tune the regularization hyperparameter so that cross-entropy was minimized. The smallest regularization parameter was chosen where cross-entropy was within one standard error of its minimum. This afforded predicting MOA of the unseen validation dataset with more than 93% accuracy across the compound classes. Finally, models were trained on the entire ground truth dataset before making predictions on unknown compounds.

**Broth microdilution assays**

The minimum inhibitory concentration of compounds was determined by making a 5 mM compound stock in DMSO which was then two-fold serially diluted ten times. Wells of a clear-bottom 96-well plate (Corning) were filled with 49 µL of appropriate medium (Middlebrook 7H9-OADC-acetate for Mycobacteria or Lysogeny Broth (LB) for *E. coli* or *P. aeruginosa*), and 1 µL compound stock was added. Finally, 50 µL exponential-phase bacterial culture which had been diluted to an $OD_{600}$ of 0.005 was added to give a final concentration series of 0.1 µM to 50µM at an initial $OD_{600}$ of 0.0025.

Plates were incubated at 37ºC in a humidified container for 24 h for *S. aureus*, *E. coli*, *P. aeruginosa*, and gram-negative species in general, 3 d for *M. smegmatis*, and 14 d for *M. tuberculosis*. Finally, $OD_{600}$ was measured using a SpectraMax M5 plate reader (Molecular Dimensions). Normalized percent inhibition (NPI) was reported using NPI = $(\mu_p - x_i) / (\mu_p - \mu_n)$, where $\mu_p$ is the mean positive control value, $\mu_n$ is the mean negative control value, and $x_i$ is the value of compound $i$. Dose response curves show an average of four independent replicates.

**DNA gyrase assays**

All compounds were screened at 160 µM in duplicate in a 96-well PCR plate (Axygen). A reaction buffer (Inspiralis) made of 50 mM HEPES-KOH (pH 7.9), 6 mM magnesium acetate, 4 mM DTT, 1 mM ATP, 100 mM potassium glutamate, 2 mM spermidine, and 0.5 mg/mL albumin was mixed with 0.5 µg relaxed pBR322 DNA (Inspiralis) for the supercoiling assay or 0.2 µg kDNA (Inspiralis) for the decatenation assay before adding test compound from a 10× stock in DMSO or water. *M. tuberculosis* DNA gyrase (Inspiralis) was diluted in 50 mM Tris-HCl at pH 7.9, 5 mM DTT, and 30 % w/v glycerol, and 2.5 U diluted DNA gyrase were added to each reaction mixture. The supercoiling and decatenation assay mixtures were incubated for 45 minutes at 37 °C. Proteinase K (Qiagen) was added to each reaction to a final concentration of 50 µg/mL and the reactions were incubated at 37 °C for an additional 30 minutes. The reactions were stopped by the addition of 20 % w/v sucrose, 50 mM Tris-HCl pH 8, 5 mM EDTA, and 0.25 mg/mL bromophenol blue. A linear pBR322 control was made via digestion of pBR322 by EcoRI-HF (NEB) for 1 h at 37 °C. DNA was extracted from the supercoiling and decatenation assay mixtures using 0.042 % v/v chloroform in isoamyl alcohol. Assay samples and the linear control were electrophoresed on a 1 % w/v agarose gel for 90 min at 80 V. The gels were stained for 15 min in 1 µg/mL EtBr, destained for 10 minutes in water, and imaged for analysis. Pixel density of bands was measured using ImageJ, and significance and log fold change was determined by a Gamma-family generalized linear model (GLM) with a log link, using DMSO untreated control as the intercept and the relaxed band density as an offset to normalize for sample loading. The two-tailed Wald test was used to determine statistical significance.

**Mycolic acid assay**
Detection of $^{14}$C-acetic acid incorporation into mycolic acids by Mtb was carried out as previously described [21]. Briefly, approximately 10 µCi $^{14}$C-acetic acid (American Radiolabeled Chemicals) was added to 10 mL exponential-phase culture with 100 µL of a 100× compound stock in DMSO to give a final concentration of 10× $MIC_{90}$ as measured by broth microdilution. After 20 h of incubation at 37 ºC with shaking, cultures were washed twice with MilliQ water, finally resuspended in 1 mL, and heat inactivated at 80 ºC for 2 h, before saponification with 1 mL aqueous 40% v/v tetrabutylammonium hydroxide at 100 ºC for 24 h. Fatty acids were methylated using 100 µL methyl iodide in 2 mL dichloromethane (DCM) at room temperature for 2 h, acidified, and extracted in DCM. Extracts were evaporated to dryness at 50 ºC for 16 h and dissolved in 200 µL DCM. Radioactivity was quantified by dilution of 50 µL sample in 5 mL scintillation fluid. Samples were normalized by counts per minute (CPM) before loading 5 µL of each on a silica thin layer chromatography (TLC) plate. Three TLC elutions of 5% v/v hexane in ethyl acetate were performed. X-ray film (Thermo Fisher) was exposed to the TLC plate at –80ºC for 2–8 d. Pixel density of bands was measured using ImageJ, and significance and log fold change was determined by a Gamma-family generalized linear model (GLM) with a log link, using DMSO untreated control as the intercept and the FAMEs band density as an offset to normalize for sample loading. The two-tailed Wald test was used to determine statistical significance.

**Metabolite supplementation assay**
Broth microdilution was performed as above, but each dose response series of a compound of interest was repeated three times, each time with a different metabolite in the growth medium: PABA at 200 µM from a 20 mM stock in ethanol; folic acid at 200 µM from a 20 mM stock in DMSO; and L-tryptophan at 1 mM from a 100 mM stock in 500 mM hydrochloric acid.

**RNA polymerase assay**
A 5′-phosphorylated, 45-nucleotide single stranded DNA (ssDNA) oligomer (Integrated DNA Technologies) was circularized using CircLigase ssDNA Ligase (Lucigen). On ice, RNA polymerase reaction buffer containing 40 mM Tris-HCl pH 7.5, 150 mM KCl, 10 mM $MgCl_2$, 1 mM DTT, and 0.01 % v/v Triton X-100 was mixed with 0.5 U of *E. coli* RNA polymerase (NEB), 8 µM DTT, 28 U RNase inhibitor (NEB), and 10 µM rifampin (positive control) or 100 µM test compound. All compounds were tested in quadruplicate. The enzyme-compound mixture

was incubated for 10 min in a 37 °C water bath. At room temperature, 2 pmol of the circularized ssDNA oligomer was added to the mixture. Reaction mixtures were transferred to 37 °C and 1 mM NTPs were added, initiating RNA transcription. The reactions were then incubated for 1 h at 37 °C. RNA was then quantified using RiboGreen fluorescent dye (Thermo Fisher). RiboGreen was diluted 1:200 in TE buffer, pH 7.5 and 100 μL of the diluted dye was mixed with 2.5 μL reaction mixture diluted with 95 μL TE buffer, pH 7.5. This mixture was incubated for 5 min at room temperature. Fluorescence was read using a plate reader with 485 nm excitation and 535 nm emission wavelengths. Significance and log fold change was determined by a Gamma-family generalized linear model (GLM) with a log link, using DMSO untreated control wells as the intercept and modeling plate-to-plate variation. The two-tailed Wald test was used to determine statistical significance.

**Generating resistant mutants**
Bacterial cultures in mid-exponential growth phase were centrifuged at 3000 × *g* for 10min and resuspended as a slurry in 1mL Middlebrook 7H9-OADC. 50 μL slurry was plated on 6 mL agar containing 2×, 4× or 8× $MIC_{90}$ of compound as determined by broth microdilution. in six-well tissue culture dishes (Corning). Dishes were incubated at 37ºC in a humidified container for more than 21 d.

At 21 d, agar was checked every 7 d for colonies. Colonies were picked into 1 mL Middlebrook 7H9-OADC-acetate in a 96-well 2 mL well volume culture block and incubated at 37 ºC for 7 d in a humidified container. Samples were then subjected to whole genome sequencing. Mutant-containing wells identified therein were used to inoculate 10 mL cultures which were grown to mid-exponential phase before storage at –80 ºC.

**Whole genome sequencing of Mycobacteria**
10 μL samples were taken from 1 mL colony-inoculated cultures and combined with 10 μL 10 % v/v DMSO in a 96-well clear round-bottom plate (Corning). Plates were heat-inactivated at 80 ºC for 2 h. Genomic DNA (gDNA) was separated from intact cells and cell debris using AMPure XP beads, eluting in 40 μL MilliQ water. Since gDNA abundance was expected to be lower than necessary for library construction, it was amplified using 6 μM random primers (Invitrogen) and φ29 DNA Polymerase (NEB) according to manufacturer's instructions using 1.5 μL gDNA and 0.2 units yeast inorganic pyrophosphatase (NEB) in 10 μL reaction volume. Reactions were incubated at 30 ºC for 24 h.

Amplified gDNA was purified using AMPure XP beads and subjected to NextEra XT (Illumina) NGS library construction according to the manufacturer's protocol. Libraries were paired-end sequenced for 150 cycles on the Illumina MiSeq platform. Reads were aligned to the AL123456 reference sequence[22] using the BWA-mem algorithm and mutations were called using the deepSNV package[23] for R.

**Efflux assay**
Efflux rates were measured as previously described [24]. Briefly, *M. smegmatis* strains were grown in Middlebrook 7H9 medium or Lysogeny Broth (LB) to an $OD_{600}$ of 0.4–0.6 and then centrifuged for 5 min at 3500 rpm. The pellet was washed once with PBS at 37 ºC and resuspended in 37 ºC PBS to give a final $OD_{600}$ of 0.4. EtBr was added to the cells at a final concentration of 0.2 μg/mL, 1 μg/mL, or 2 μg/mL, and bacteria were incubated for 30 minutes at 37 °C. After EtBr treatment, cells were centrifuged for 5 min at 3500 rpm and resuspended in 37

°C PBS to give a final $OD_{600}$ of 0.8. A white 96-well plate (Corning) was prepared with serially diluted compound and 50µL PBS containing 0.8% w/v glucose. 50µL EtBr-loaded bacteria were added to each well of the plate. Fluorescence was read at 37 °C in a SpectraMax M5 (Molecular Dimensions) plate reader using 530 nm excitation and 585 nm emission wavelengths and was recorded every 30 s for 2 h.

An exponential decay model ($F(t) = b + F(0)\exp(-k_{obs}t)$, where $F(t)$ is fluorescence at time $t$, $b$ is the baseline fluorescence, and $k_{obs} = k_{in} + k_{out}$ is the exponential decay rate) was fitted to the trace for each well in the experiment. A non-competitive inhibition model ($k_{obs} = a + c / (1 + IC_{50} / [I])$, where $IC_{50}$ is the inhibitor dissociation constant, and [I] is inhibitor concentration; $a$ and $c$ are baseline and amplitude parameters) was fitted to the $k_{obs}$ dependence on compound concentration. Models were fitted by non-linear least squares using the nls command in R.

**Data availability**

The raw primary screen data and calculated log fold changes and *p*-values are available online at https://broad.io/cgtb.

**Code availability**

ConcensusGLM is available at http://github.com/eachanjohnson/concensusGLM. Other computer code is available from the corresponding author upon request.


1 Kim, J. H. *et al.* Protein inactivation in mycobacteria by controlled proteolysis and its application to deplete the beta subunit of RNA polymerase. *Nucleic Acids Res.* **39**, 2210-2220, doi:10.1093/nar/gkq1149 (2011).

2 Murphy, K. C., Papavinasasundaram, K. & Sassetti, C. M. Mycobacterial recombineering. *Methods Mol. Biol.* **1285**, 177-199, doi:10.1007/978-1-4939-2450-9_10 (2015).

3 Grant, S. S. *et al.* Identification of novel inhibitors of nonreplicating *Mycobacterium tuberculosis* using a carbon starvation model. *ACS Chem. Biol.* **8**, 2224-2234, doi:10.1021/cb4004817 (2013).

4 Wellington, S. *et al.* A small-molecule allosteric inhibitor of *Mycobacterium tuberculosis* tryptophan synthase. *Nat. Chem. Biol.* **13**, 943-950, doi:10.1038/nchembio.2420 (2017).

5 Stanley, S. A. *et al.* Identification of novel inhibitors of *M. tuberculosis* growth using whole cell based high-throughput screening. *ACS Chem. Biol.* **7**, 1377-1384, doi:10.1021/cb300151m (2012).

6 Reynolds, R. C. *et al.* High throughput screening of a library based on kinase inhibitor scaffolds against *Mycobacterium tuberculosis* H37Rv. *Tuberculosis (Edinb)* **92**, 72-83, doi:10.1016/j.tube.2011.05.005 (2012).

7 Ananthan, S. *et al.* High-throughput screening for inhibitors of *Mycobacterium tuberculosis* H37Rv. *Tuberculosis (Edinb)* **89**, 334-353, doi:10.1016/j.tube.2009.05.008 (2009).

8 Maddry, J. A. *et al.* Antituberculosis activity of the molecular libraries screening center network library. *Tuberculosis (Edinb)* **89**, 354-363, doi:10.1016/j.tube.2009.07.006 (2009).

9 Ballell, L. *et al.* Fueling open-source drug discovery: 177 small-molecule leads against tuberculosis. *ChemMedChem* **8**, 313-321, doi:10.1002/cmdc.201200428 (2013).

10 Chou, D. H. *et al.* Synthesis of a novel suppressor of beta-cell apoptosis via diversity-oriented synthesis. *ACS Med. Chem. Lett.* **2**, 698-702, doi:10.1021/ml200120m (2011).
11 Love, M. I., Huber, W. & Anders, S. Moderated estimation of fold change and dispersion for RNA-seq data with DESeq2. *Genome Biol.* **15**, 550, doi:10.1186/s13059-014-0550-8 (2014).
12 McCarthy, D. J., Chen, Y. & Smyth, G. K. Differential expression analysis of multifactor RNA-Seq experiments with respect to biological variation. *Nucleic Acids Res.* **40**, 4288-4297, doi:10.1093/nar/gks042 (2012).
13 R: A language and environment for statistical computing (R Foundation for Statistical Computing, Vienna, Austria, 2017).
14 Venables, W. N. & Ripley, B. D. *Modern Applied Statistics with S*. Fourth edn, (Springer, 2002).
15 Kanehisa, M., Furumichi, M., Tanabe, M., Sato, Y. & Morishima, K. KEGG: new perspectives on genomes, pathways, diseases and drugs. *Nucleic Acids Res.* **45**, D353-D361, doi:10.1093/nar/gkw1092 (2017).
16 van der Maaten, L. & Hinton, G. Visualizing Data using t-SNE. *Journal of Machine Learning Research* **9**, 2579-2605 (2008).
17 Rtsne: t-Distributed Stochastic Neighbor Embedding using a Barnes-Hut Implementation (2015).
18 Murtagh, F. & Legendre, P. Ward's Hierarchical Agglomerative Clustering Method: Which Algorithms Implement Ward's Criterion? *Journal of Classification* **31**, 274-295, doi:10.1007/s00357-014-9161-z (2014).
19 Tibshirani, R., Walther, G. & Hastie, T. Estimating the number of clusters in a data set via the gap statistic. *J. Roy. Stat. Soc. Ser. B. (Stat. Method.)* **63**, 411-423 (2001).
20 Friedman, J., Hastie, T. & Tibshirani, R. Regularization Paths for Generalized Linear Models. *Journal of Statistical Software* **33**, 1-22 (2010).
21 Vilcheze, C. & Jacobs, W. R. Isolation and Analysis of *Mycobacterium tuberculosis* Mycolic Acids. *Current Protocols in Microbiology* **5**, 10A.13.11-10A.12.11, doi:10.1002/9780471729259.mc10a03s05 (2007).
22 Cole, S. T. *et al.* Deciphering the biology of *Mycobacterium tuberculosis* from the complete genome sequence. *Nature* **393**, 537-544, doi:10.1038/31159 (1998).
23 Gerstung, M. *et al.* Reliable detection of subclonal single-nucleotide variants in tumour cell populations. *Nat Commun* **3**, 811, doi:10.1038/ncomms1814 (2012).
24 Paixao, L. *et al.* Fluorometric determination of ethidium bromide efflux kinetics in *Escherichia coli*. *J. Biol. Eng.* **3**, 18, doi:10.1186/1754-1611-3-18 (2009).

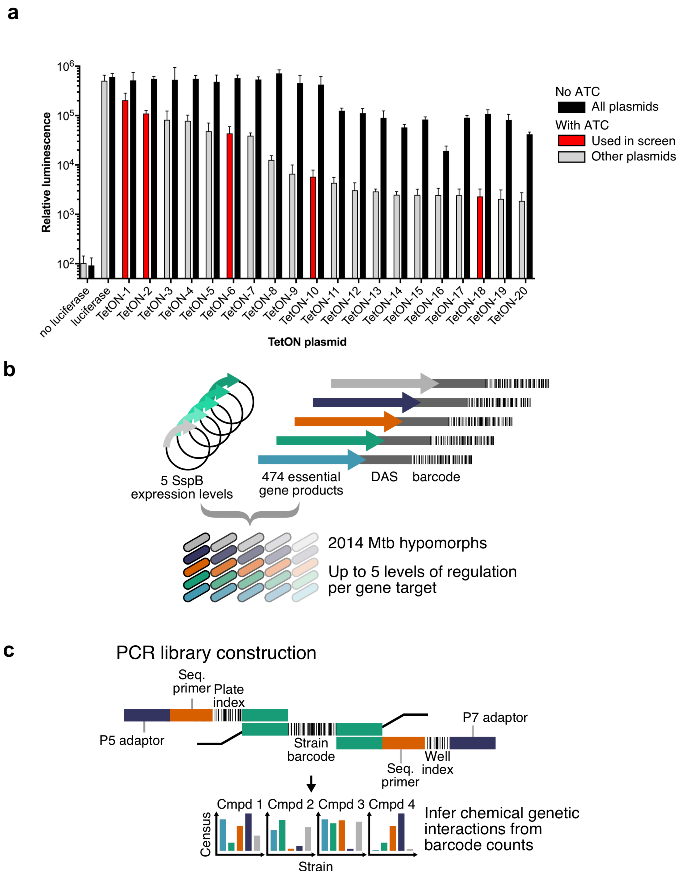


**Extended Data Figure 1 | Mtb hypomorph strain creation.**

**a**, Degradation of a DAS-tagged target gene product was mediated by SspB, whose expression was driven by an ATC-inducible TetON promoter. To allow individualized degrees of knockdown for each gene product, a series of TetON promoters with varying strengths was generated. Regulated promoter strength was quantified by fusion to a luciferase gene and measuring luminescence in the presence and absence of the ATC inducer. In the screen, strains containing the TetON-2, -6, -10, and -18 promoters were used for subsequent strain construction.
**b**, A range of up to five different knock-down levels were attempted using five different promoters for each target gene, allowing generation of 2014 hypomorphs.
**c**, Chromosomal strain barcodes were inserted into each engineered hypomorph, thus allowing PCR amplification by an array of primers containing 5′-overhangs encoding screen location (well and plate) barcodes. For census enumeration of pooled strains, PCR products were combined and subjected to Illumina NGS.

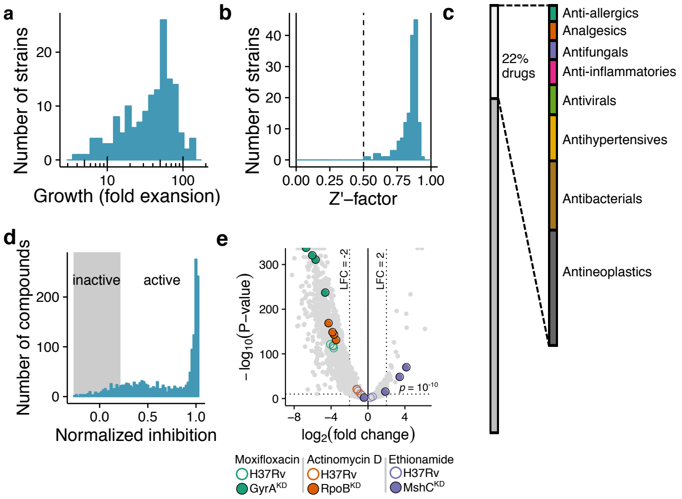


**Extended Data Figure 2 | Strain and compound collection characterization.**
**a**, Histogram of growth of the individual hypomorphs in the screening pool over the 14-day duration of the screen. Most hypomorphs were within a 10-fold window of growth rates.
**b**, Histogram of hypomorph Z′-factors between the untreated wells and rifampin control wells of the screening assay for the hypomorphs. All Z′-factors were greater than 0.5, indicating an excellent screening assay.
**c**, The composition of the bioactive compound library. The right bar shows the broad classes within the segment of known drugs.
**d**, The bioactive compound library was tested up to 50 µM against a GFP-expressing Mtb strain. Most compounds had detectable activity for at least one concentration tested.
**e**, Volcano plot of chemical-genetic interactions from the bioactive library. Each point represents a single strain-compound interaction at a single concentration. Some interactions of interest are highlighted: compounds are designated by color, while wild-type Mtb interactions are shown in open circles and hypomorphs interactions of interest are shown in solid circles. The majority of interactions were inhibitory, since the compound library was confirmed to be enriched for antitubercular activity.

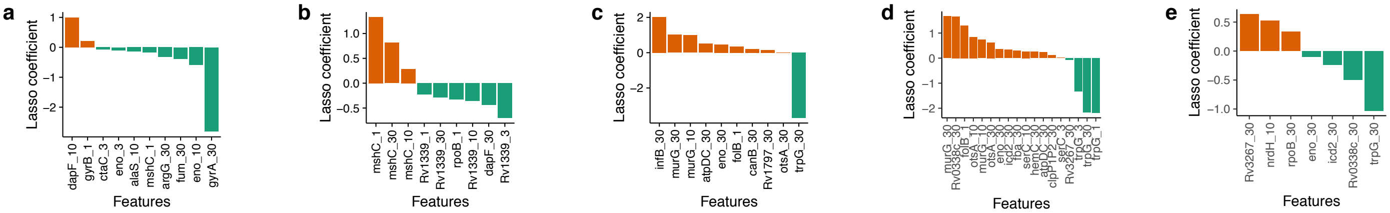


**Extended Data Figure 3 | Feature weights of Lasso classifiers.**

**a**, Feature weights of the hypomorphs used by the Lasso binary classifier to predict DNA gyrase inhibitors. We trained on the fluoroquinolones, with the GyrA hypomorph being the most predictive strain. Features are denoted as hypomorph's target gene and compound concentration separated by an underscore.

**b**, As (a) for the Lasso binary classifier to predict mycolic acid biosynthesis inhibitors. We trained on known InhA inhibitors, with the MshC hypomorph being a prominent discriminator.

**c**, As (a) for the Lasso binary classifier to predict folate biosynthesis inhibitors. We trained on the sulfonamides, with the TrpG hypomorph being a prominent discriminator.

**d**, As (a) but trained on chemical-genetic interaction profiles from the sulfonamides and confirmed new folate biosynthesis inhibitors.

**e**, As (a) but trained on chemical-genetic interaction profiles of the newly-identified tryptophan biosynthesis inhibitors.

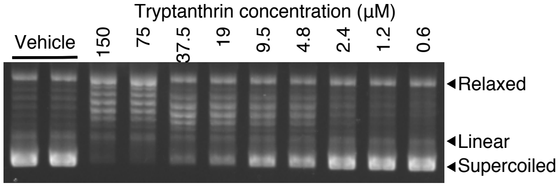


**Extended Data Figure 4 | DNA gyrase inhibition by tryptanthrin.**

Agarose gel showing increasing inhibition of Mtb DNA gyrase supercoiling activity with increasing tryptanthrin concentration. DNA gyrase catalyzes supercoiling of pBR322; inhibitors prevent the accumulation of supercoiled gel bands.

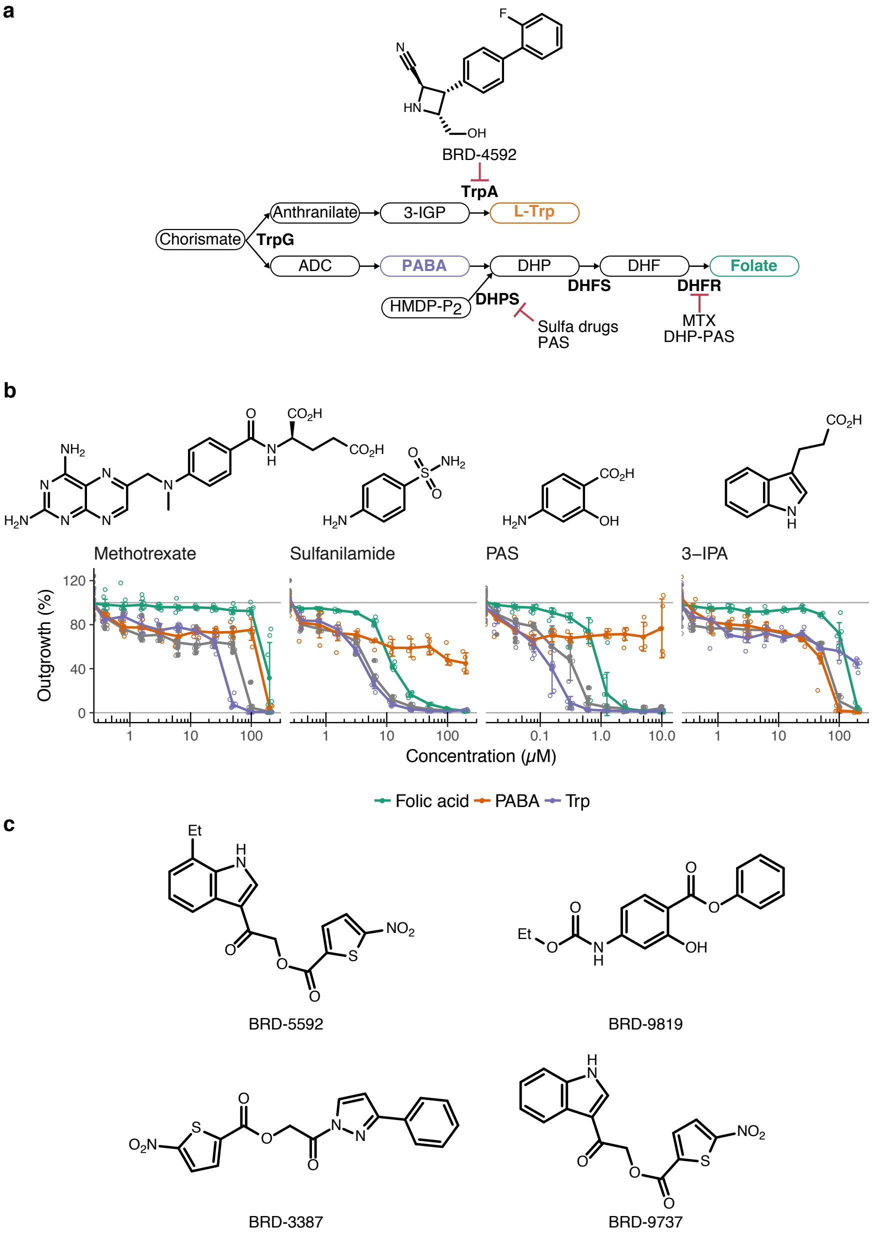


**Extended Data Figure 5 | New classes of inhibitors of folate and tryptophan biosynthesis.**
**a**, Schematic of the folate and tryptophan biosynthesis pathways. TrpG is an amphibolic enzyme, upstream of both PABA and tryptophan. Biosynthetic enzymes mentioned in the text are indicated in their metabolic context. 3-IGP: 3-indoleglycerol phosphate; ADC: 4-amino-4-deoxychorismate; PABA: para-amino-benzoic acid; DHP: dihydropteroate; DHF: dihydrofolate; HMDP-$P_2$: 6-hydroxymethyl-7,8-dihydropterin diphosphate; DHPS: dihydropteroate synthase; DHFS: dihydrofolate synthase; DHFR: dihydrofolate reductase; Sulfa drugs: sulfonamide antibiotics; MTX: methotrexate; PAS: *para*-aminosalicylic acid; DHP-PAS: adduct of DHP and PAS.
**b**, Dose response curves of known folate biosynthesis inhibitors and the validated tryptophan biosynthesis inhibitor scaffold, 3-indole propionic acid, supplemented with either PABA, folic

acid, or tryptophan. Chemical structures of the known inhibitors are shown. Individual replicates ($n$ = 4) are shown as open circles, means are shown as filled circles, and error bars show 95% confidence intervals.
**c**, Chemical structures of predicted and subsequently validated folate biosynthesis inhibitors, including the nitrothiophenes and *para*-aminosalicylic acid derivative BRD-9819.

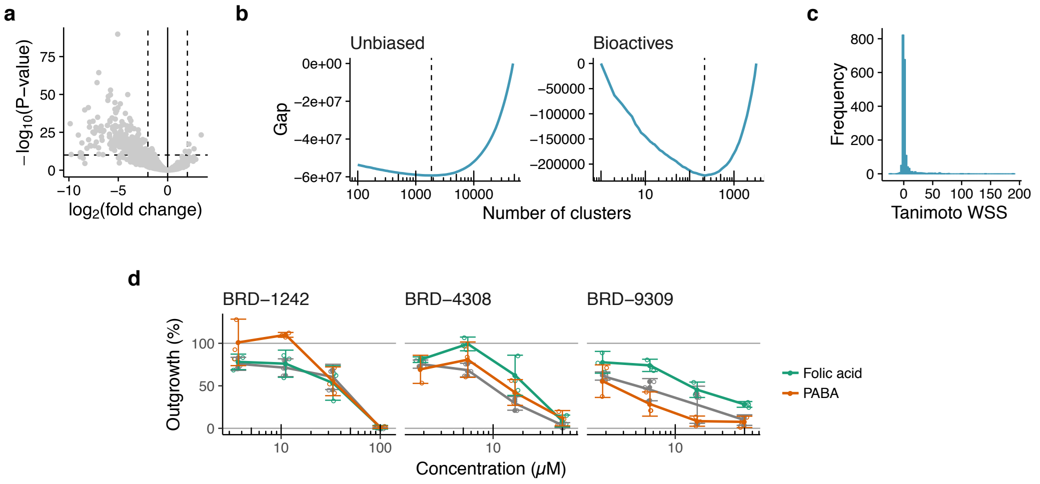


**Extended Data Figure 6 | Performance of a large, unbiased compound library.**
**a**, Volcano plot of chemical-genetic interactions from the larger unbiased chemical library. Each point represents a single strain-compound interaction at a single concentration.
**b,** Clustering of chemical-genetic interaction profiles. The number of chemical-genetic interaction profile clusters in the two libraries was determined by finding the minimum Gap statistic, a measure of within-cluster similarity compared to clustering at random. The minimum, denoted by the dotted lines, shows the estimate of the true number of clusters in the unbiased and bioactive libraries, with the unbiased library containing many more unique chemical-genetic clusters.
**c,** Chemical-genetic clusters are enriched for chemically similar compounds in the unbiased library. The *y*-axis shows the frequency of chemical-genetic clusters with a particular Tanimoto WSS Z-score (*x*-axis), which is an indicator of in-cluster chemical similarity. One-third of the clusters have meaningful structure-activity relationships, that is, compounds within a cluster have significantly greater chemical similarities than by chance.
**d**, Actual compound performance of predicted folate biosynthesis inhibitors in a metabolite rescue assay. Mtb was treated with predicted inhibitors in the presence or absence of folate or PABA. The effect of BRD-1242 is abolished by PABA alone, while the effects of BRD-4308 and BRD-9309 are abolished by folate alone, suggesting that they are inhibitors of folate biosynthesis with distinct mechanisms. Individual replicates ($n$ = 3) are shown as open circles, means are shown as filled circles, and error bars show 95% confidence intervals.

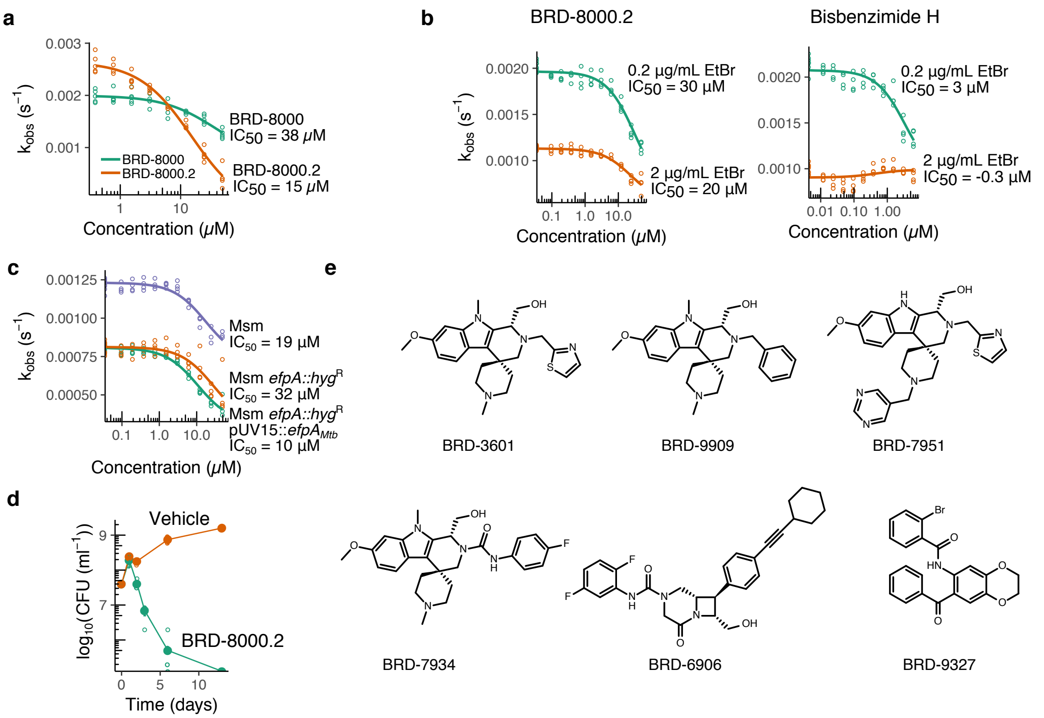


**Extended Data Figure 7 | Inhibitors of a new target, EfpA, in *M. tuberculosis*.**

**a**, As in (a), but for the initial hit BRD-8000 and the whole cell activity optimized derivative, BRD-8000.2. The relative rate constants correlate with their potency against wild type Mtb. Individual replicates are shown as open circles, and fitted curves are shown as lines.

**b**, BRD-8000.2 inhibition of efflux is independent of EtBr concentration. Measurements of first-order rate constant for EtBr efflux, $k_{\mathrm{obs}}$, against varying concentration of BRD-8000.2 or bisbenzimide H and two concentrations of the EtBr efflux substrate. BRD-8000.2 is a non- or an un-competitive inhibitor in contrast to bisbenzimide H, a known competitive efflux inhibitor.

**c**, As in (a), but with BRD-8000.2 and wild type Msm, an *efpA*-knockout strain of Msm, and the knockout strain complemented with $efpA_{Mtb}$. The higher $IC_{50}$ for the knockout strain shows that BRD-8000 efflux inhibition is EfpA-specific; complementation restores the lower $IC_{50}$.

**d**, BRD-8000.2 is bactericidal as demonstrated by colony forming unit enumeration over time. Individual replicates ($n$ = 8) are shown as open circles, means are shown as filled circles, and error bars show 95% confidence intervals.

**e**, Additional compounds predicted and validated to be efflux inhibitors.
